# Single-cell immuno-mechanics: rapid viscoelastic changes are a hallmark of early leukocyte activation

**DOI:** 10.1101/851634

**Authors:** Alexandra Zak, Sara Violeta Merino Cortés, Anaïs Sadoun, Avin Babataheri, Stéphanie Dogniaux, Sophie Dupré-Crochet, Elodie Hudik, Hai-Tao He, Abdul I Barakat, Yolanda R Carrasco, Yannick Hamon, Pierre-Henri Puech, Claire Hivroz, Oliver Nüsse, Julien Husson

## Abstract

To accomplish their critical task of removing infected cells and fighting pathogens, leukocytes activate by forming specialized interfaces with other cells. Using an innovative micropipette rheometer, we show in three different cell types that when stimulated by microbeads mimicking target cells, leukocytes become up to ten times stiffer and more viscous. These mechanical changes initiate within seconds after contact and evolve rapidly over minutes. Remarkably, leukocyte elastic and viscous properties evolve in parallel, preserving a well-defined ratio that constitutes a mechanical signature specific to each cell type. The current results indicate that simultaneously tracking both elastic and viscous properties during an active cell process provides a new way to investigate cell mechanical processes. Our findings also suggest that dynamic immuno-mechanical measurements provide an identifier of leukocyte type and an indicator of the cell’s state of activation.

## Introduction

The understanding of cell mechanics has progressed thanks to the development of micromanipulation techniques such as micropipette aspiration^1–9^, indentation techniques^10–18^, and others^19–21^. Microfluidics-based approaches now allow high-throughput mechanical measurements^22–29^. Yet, these techniques make it difficult to track mechanical changes in cells stimulated by soluble molecules^30,31^ or activating surfaces^32,33^. Tracking these changes can bring a wealth of information on cell function and disease.

Here we focus on white blood cells (leukocytes), which – among several other functions – fight dead and infected cells or pathogens. To do so, leukocytes activate by forming specialized interfaces called synapses with other cells. Different leukocytes form different types of synapses, which share several molecular^34^ but also mechanical features. For instance, we have shown that when forming a synapse, a T lymphocyte generates forces^35,36^ and stiffens^37^, recapitulating pioneering observations that during phagocytosis, the cortical tension of a phagocytic leukocyte increases dramatically while the cell engulfs its prey^6,38–40^. Prior to the work presented here, the mechanics of other leukocytes (such as B lymphocytes) remained unknown, and some common mechanical aspects such as the evolution of viscous properties of leukocytes during activation were to date unexplored. Cellular matter is indeed particular in that elastic and viscous properties of cells are usually *not* independent: resting leukocytes such as neutrophils^41,42^ and macrophages ^6^ conserve a ratio of cortical tension to cell viscosity of about 0.2-0.3 µm/s, even though macrophages can be ten times more tense than neutrophils. Many studies showed this peculiar aspect of biological matter in other cell types^20,43–48^. What remained unknown was whether this link between elastic and viscous properties would be conserved over time even during mechanical changes due to cell function, e.g. activation.

In this study, we quantified the evolution of both elastic and viscous properties during the activation of three types of leukocytes, and addressed the evolution of both elastic and viscous properties over time. We used a micropipette rheometer to activate leukocytes with standardized activating anti-body-covered microbeads^37,49,50^. Aside we probed early mechanical changes occurring in leukocyte-target cell contacts by using Atomic Force Microscopy.

## Results

### Micropipette rheometer for monitoring rapid morphological and mechanical changes in non-adherent cells

We implemented a real-time feedback loop in our Micropipette Force Probe setup^37,49^ allowing us to impose a controlled small oscillatory force modulation Δ*F* cos(ω*t*) (angular frequency ω = 2π*f*, frequency *f* = 1 Hz) superimposed onto a constant force 〈*F*〉. A total force *F*(*t*) = 〈*F*〉 + Δ*F* cos(ω*t*) is thus applied to the leukocyte during its activation following the contact with an activating microbead. ensured by pressing the bead against the cell with a force *F_comp_* (0.12-0.36 nN). This initial compression, from which we extract the cell's effective Young’s modulus *E_Young_*^51,52^ (Suppl. Mat. 1), is followed by an imposed force modulation regime, where we measure oscillations in the position of the tip of the flexible micropipette *x_tip_* = 〈*x_tip_*〉 + Δ*x_tip_* cos(ω*t* − φ), of average value 〈*x_tip_*〉, amplitude Δ*x_tip_*, (typically 100 nm, with a few-nanometer accuracy) and phase lag φ. Changes in *x_tip_* reflect changes in cell length (Figure 1). From *x_tip_* and *F*(*t*), we deduce the complex cell stiffness *K*^∗^ = *K*^′^ + *iK*^″^, of elastic part *K’* and viscous part *K”* (Suppl. Mat. 2, Suppl. Movie S1). We validated our setup on red blood cells (suppl. movie S2), for which existing models predict *K’* (Suppl. Mat. 3).

**Figure 1.**
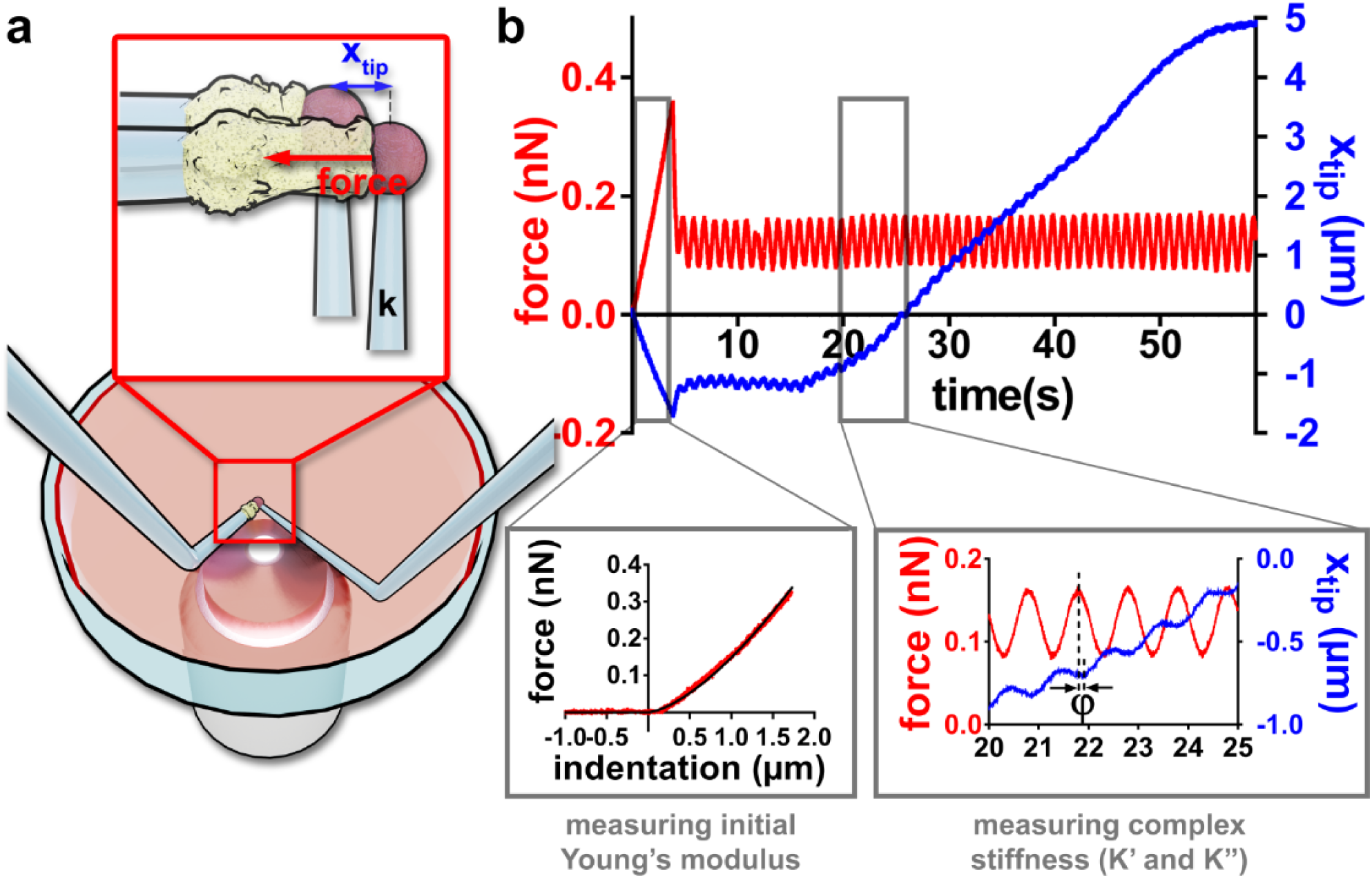
Rheology during leukocyte activation. (**a**). Setup. Two micropipettes are plunged in a Petri dish. A flexible pipette (right, bending stiffness *k*≈0.2 nN/µm) holds an activating microbead firmly. A rigid micropipette (left) gently holds a leukocyte (aspiration pressure of 10-100 Pa). The base of the flexible micropipette is displaced to impose a desired force on the cell. (**b**) Force applied on the cell (in red) and the position *x_tip_* of the tip of the flexible micropipette (in blue). The cell is first compressed with a force *F_comp_* (here 0.36 nN) to measure the initial Young’s modulus of the cell (inset, bottom left). Then an oscillatory force is applied to the cell, leading to an oscillatory *x_tip_* signal (inset, bottom right) superimposed to a slower variation of the average value <*x_tip_*> due to the growth of a protrusion produced by the leukocyte.

### Viscoelastic changes during leukocyte activation

We probed three different types of leukocytes during activation and observed qualitatively similar behavior: both *K’* and *K”* increase and reach a maximum that is 3- to 8-fold larger than their initial value within few minutes (~2 minutes for T and B cells, ~5 minutes for neutrophils-like PLB985 cells) following the contact between the leukocyte and the relevant activating microbead (Figure 2, Suppl. Movie S1).

**Figure 2.**
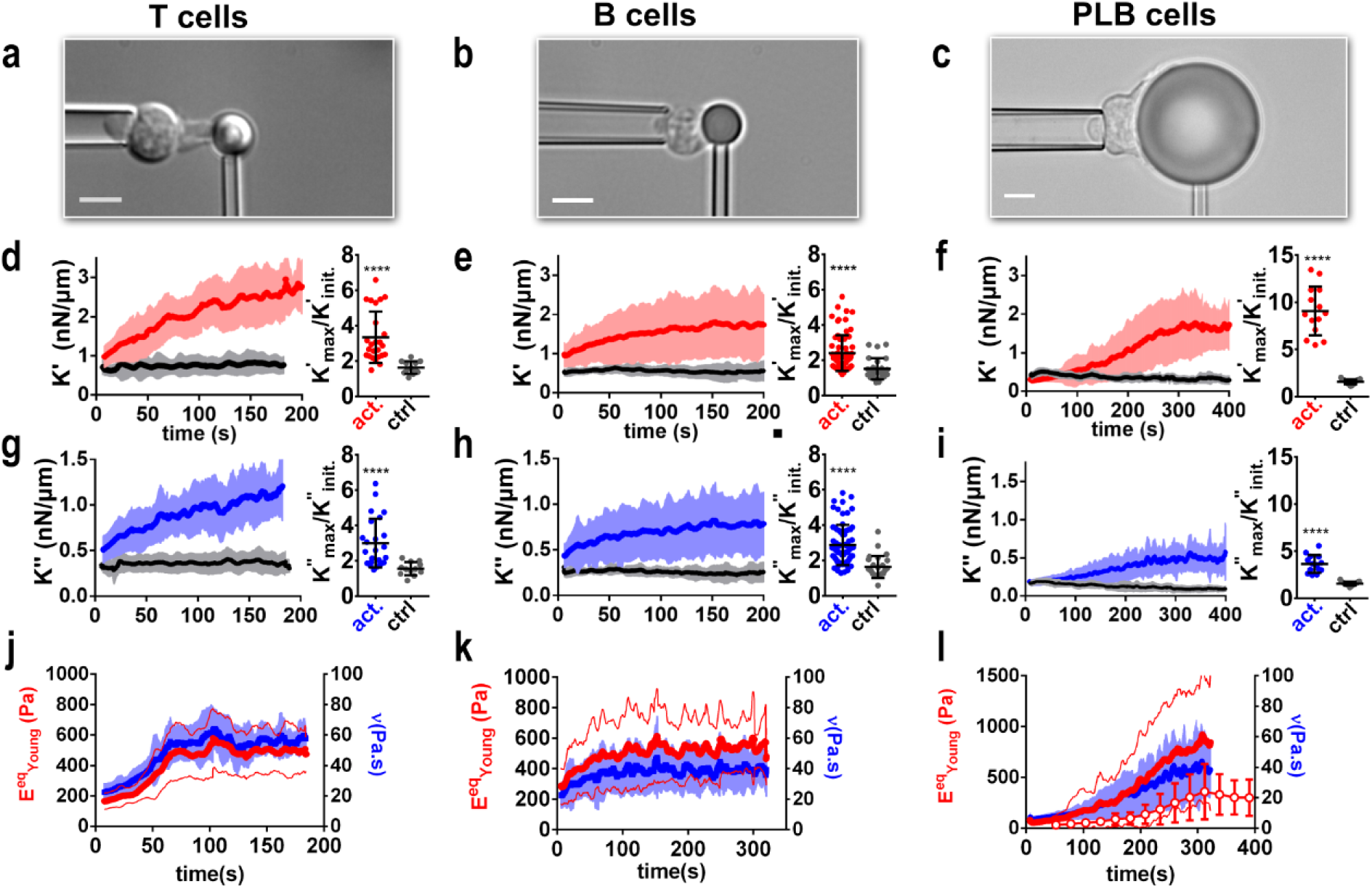
Both *K’* and *K”* increase during leukocyte activation. Each column corresponds to a different cell type. (**a-c**) Cell morphology. Scale bars are 5 µm. (**d-f**) Left: *K’* for activating (red lines) and non-activating control beads (gray lines). Right: max/min ratio of *K’* values, for activating and non-activating control beads. (**g-i**) Left: values of *K”* corresponding to K’ values in **d-f**. (**j-l**) Equivalent Young’s modulus (red, left axis) and cell viscosity (blue, right axis). In **l,** dots are Young’s modulus measured in the back of the cell. In all panels, leukocyte-bead contact time (t=0) is detected as a force increase, thick lines are mean±SD over at least 3 pooled experiments (T cells: 21 cells, B cells 71 cells, PLB cells: 14 cells). Mann-Whitney test, ****: p<0.0001.

### Changes in *K’* and *K”* correspond to changes in intrinsic leukocyte mechanical properties

Both *K’* and *K”* depend on cell geometry, so to exclude the possibility that changes in *K’* and *K”* only reflect geometrical changes, cell intrinsic mechanical properties such as Young’s modulus and viscosity have to be extracted from *K’* and *K”*. To do so, we first convert K’ and K” into a storage modulus E’ and a loss modulus E”, respectively. In the case of T cells and B cells, we do so by modeling cell geometry (Suppl. Mat. 5). For PLB cells, we use a modified setup where a non-adherent glass bead indents the cell on its “back” (figure 3a)^16^ while the cell phagocytoses an activating bead at its “front” (Fig. 3a, Suppl. Movie S3). This alternative setup leads to the same increase in *K’* and *K”* as measured using the original setup with front indentation (Fig. 3b). Having a sphere-against-sphere geometry, we can use a linearized Hertz model to extract *E’* and *E”* moduli (Suppl. Mat. 4). Lastly, we performed cyclic indentation experiments to measure directly the Young’s modulus over time (with a limited time resolution of ~30 seconds, dotted curve in Fig. 2l): indentation occurs only for a short time at the beginning of each cycle, after which the indenter retracts from the cell, marking the end of a cycle, immediately followed by a new cycle (Suppl. Movie S4). For the sake of a simple interpretation of *E’* and *E”*, which are obtained in a regime of oscillatory forces, we convert them respectively into an effective Young’s modulus 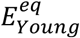 and an effective viscosity *ν*. To calculate 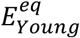, we compare the Young’s modulus measured by initial compression and *E’* measured right after, when force modulation begins (Suppl. Mat. 5; Suppl. Fig. S2). For PLB cells, we used again cyclic back-indentation described above to measure both Young’s modulus and *E’* not only at initial time but during the whole activation. This showed that *E*^′^ and *E_young_* are proportional, with a phenomenological coefficient *C* such that *E*^′^ = *CE_young_*. *C* is constant over time for PLB cells, and for T and B cells we assume that *C* is also constant over time (Suppl. Mat. 5). Using this approach, we measured the effective Young’s modulus of the three cell types over time (Fig. 2j-l). To calculate the effective cell viscosity, we consider a simple model consistent with a newtonian liquid and cortical tension model of a cell^18^, where a Newtonian viscosity is introduced as *υ* = *E*^″^⁄ω.

**Figure 3.**
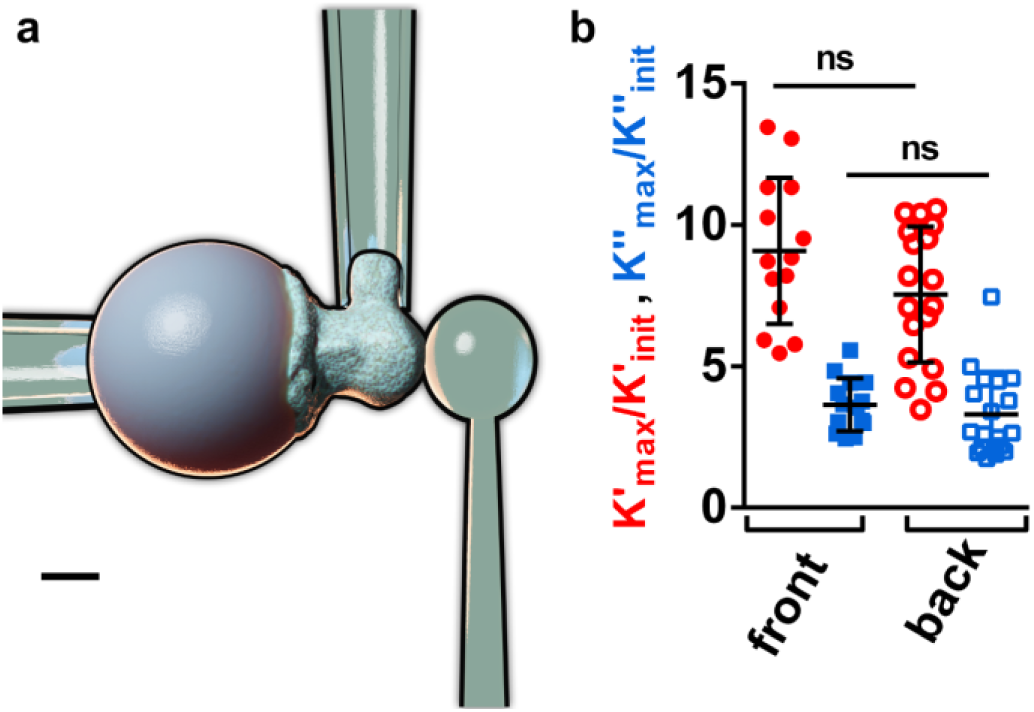
Indenting the “back” of a phagocyte during activation. (**a**) Modified setup. A stiff pipette holds the activating microbead. Scale bar is 5 µm.= (left), an auxiliary (stiff) pipette holds the cell by its “side” (top) and a flexible pipette whose tip consists of a non-adherent glass bead indents the cell on its “back” (right). (b) Max/min ratios for *K’* and *K”* obtained with both versions of the setup during PLB cell activation (ns: non-significant, two-tailed unpaired t test).

The resulting Young’s modulus and viscosity increase until reaching a maximal value that is 2-3 folds higher than the baseline value for T and B cells, and about 11 folds higher for PLB cells (Fig. 2j-l).

### Localization of mechanical changes

We asked if mechanical changes were localized close to the activation area in the leukocyte. In the case of T cells, we used a modified setup similar to the one allowing to indent PLB cells on their back, and showed that cell stiffening is higher in the vicinity of the contact zone with the activating surface (Suppl. Mat. 6, Suppl. Movie S5).

### Tension *vs. K’*

Mechanical changes precede morphological ones. Indeed, in T cells, *K’* increases slowly within seconds after cell-bead contact, before cell morphology starts changing (i.e. before the T cell produces a large protrusion^37^, Suppl. Movie S1). The onset of this growth is followed by a short period during which *K’* stays relatively constant. This phase ends by a fast increase of *K’* during protrusion growth (Fig. 4). Of note, T cells have large surface membrane stores, so it is unlikely that the acceleration in *K’* could be explained by limited membrane stores (Suppl. Mat. 7). One should also note that when *K’* starts increasing more rapidly, the tail of the cell in the holding micropipette starts retracting (Suppl. Fig. S3a), as a consequence of an increase in cell tension. We can (i) estimate a cell tension from the moment of retraction of the cell tail and (ii) compare it to an equivalent cell tension derived from *K’* based on physical considerations, and they are in good agreement (Suppl. Mat. 8) –*K’* and tension are two ways to describe cell mechanical properties.

**Figure 4.**
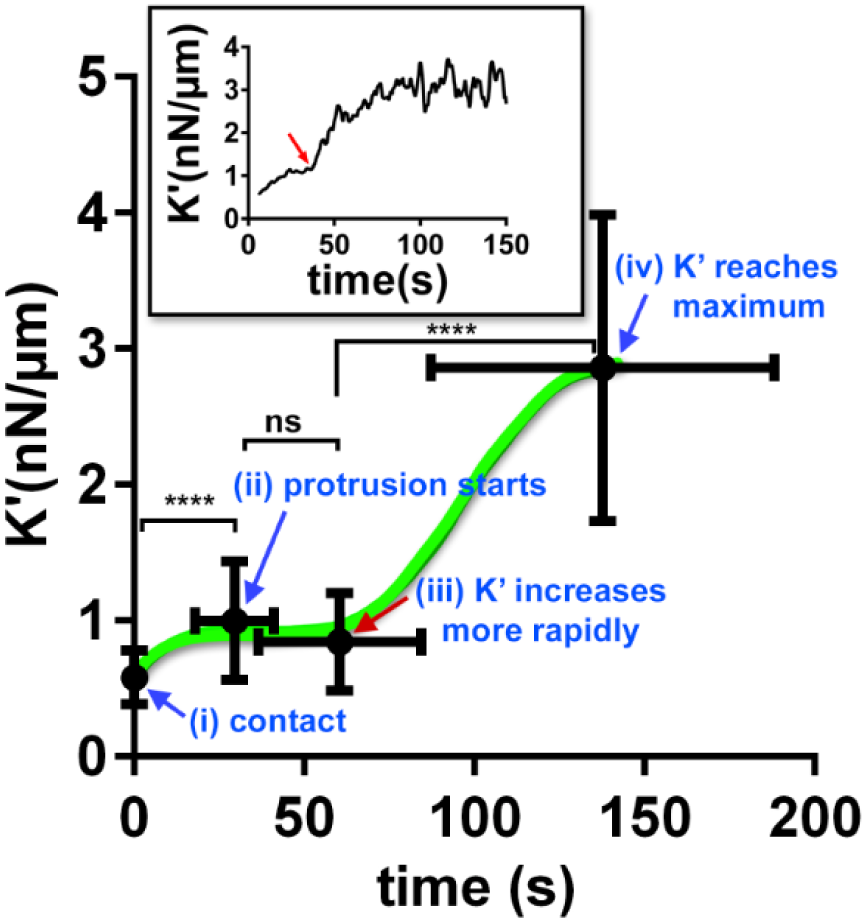
*K’* and morphology evolution during T cell activation: (i) cell-bead contact (origin of time), (ii) beginning of the growth of a protrusion produced by the cell^37^, (iii) onset of more rapid increase in *K’*, (iv) reaching maximal stiffness. Green line is only a guide to the eye. Black dots are mean+/-SD (n=18 to 39 cells over at least 3 pooled experiments). **Inset**: individual example, the red arrow indicates when *K’* increases more rapidly. Tests on K’ values are two-tailed Wilcoxon matched-pairs signed rank test, ****: p<0.0001).

### AFM measurements of early mechanical changes following T cell/APC contact

In order to confirm that T cell mechanics may be modulated within seconds when it encounters an APC before any morphological change occurs, and with an independent technique, we used AFM and performed Single Cell Force Spectroscopy (SCFS) experiments (Fig. 5a, Suppl. Mat. 9)^53^ on murine anti-CD45-immobilized 3A9m T cells encountering COS-7 surrogate antigen-presenting cells (APCs)^54,55^. To measure rapid mechanical changes using this technique, we prevented potential large active cell deformations by using the pan-Src kinase inhibitor PP2 (methods) to block the initiation of TCR-dependent signaling cascades. The contact mechanics at initial time were not modified by the presence or absence of the HEL (Hen Egg Lysozyme, peptide p46.61)-derived peptide on the APC (Supp. Fig. S6d). However, the peptide induced a larger detachment force and more numerous detachment events, indicating that the interaction is indeed peptide specific (Suppl. Fig. S6e-f).

**Figure 5.**
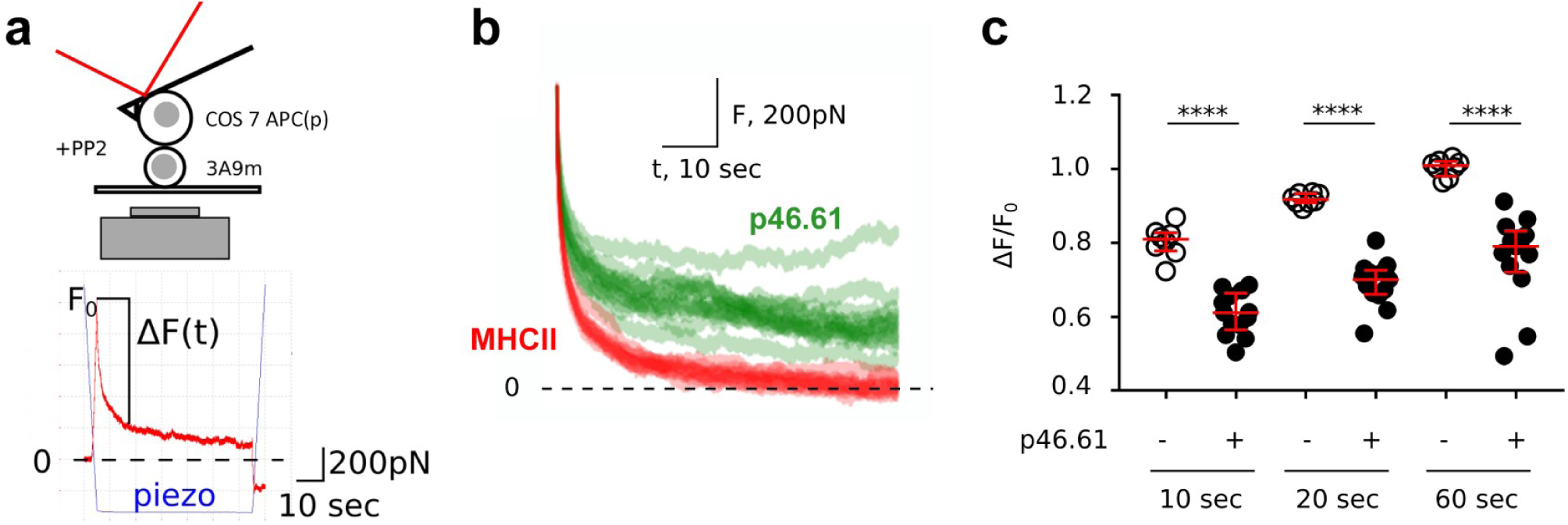
Cell-cell presentation induces early changes in force relaxation. (**a**) AFM SCFC Setup (top) and example of individual force vs. time curve (bottom). The piezo signal is the position of the base of the cantilever. (**b**) Relaxation part of force curves, for peptide (p46.61) *vs.* no-peptide (MHCII) cases. One curve corresponds to one cycle, ie. 1 APC/T cell couple. See suppl. fig. S6g for plot of mean±SD curves. (**c**) Quantification of the relaxation for three different time points, for MHCII (-) and MHCII/p46.61 (+) cases. ΔF and F_0_ definition as shown in **a** (**** for p<0.001).

We observed a striking difference in the mechanics of the cell conjugates in presence vs. absence of a saturating concentration of antigenic peptide as measured by the force relaxation after pressing a cell against the other (Fig. 5b). T cell/APC contact thus induced mechanical changes, within few seconds, and without any TCR signaling. Furthermore, the smaller amplitude in force relaxation *vs.* time due to T cell-APC contact points toward a more viscous behavior of cells following peptide recognition at the interface between a T cell and an APC, qualitatively consistent with our micropipette experiments.

### Relationship between elastic and viscous properties during leukocyte activation

We asked if variations in elastic and viscous properties were related to each other. Prior works show that in several cell types the loss modulus *E*^″^ and storage modulus *E*^′^ of a cell are related to each other. In fact their ratio 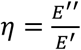, the loss tangent, lies within a narrow range of 0.1-0.3^20,30,45,46,56–59^. This implies that the stiffer the cell, the more viscous it is. This narrow range of loss tangent is consistent with the conserved ratio of cortical tension to cell viscosity in leukocytes (see introduction). In biological matter, similar to soft glassy materials ^60,61^, stress relaxation over time often follows a power-law (with an exponent that we shall call α)^43,45–48^. The loss tangent as defined by the ratio of loss and storage moduli is frequency-dependent, but is expected to be approximately equal to 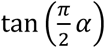 at low frequencies (Suppl. Mat. 10). Therefore, by measuring the power-law exponent α independently, we should recover the value of 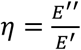 (which is also equal to 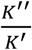 as the contribution from geometry is eliminated in this ratio^31^). To test this prediction, we performed stress relaxation experiment on resting PLB cells (fig. 6c) and obtained values for α very consistent with the predicted value of 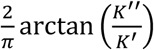 (which is equivalent to comparing 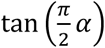 and 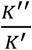, dark green star symbol in Figure 6b). We then compared *K” vs. K’* curves obtained during activation of T cells, B cells, and PLB cells (figure 6a). In Figure 8b we plot the corresponding value of 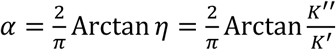: these curves were remarkably consistent for T and B cells; for PLB cells, *K”-K’* curves obtained by indentation in the front and in the back of the cell are very similar, and differ from the T and B cell common curve. Interestingly, Bufi *et al.*^62^ measured storage and loss moduli of various leukocytes at resting state or after priming by different inflammatory signals. Their data lead to a loss tangent η consistent with our measurements (Fig. 6d). Their measurements are made at a single time point only, while Maksym et al.^30^ tracked over time both the loss and storage moduli of human airway smooth muscle cells contracting due to administration of histamine. Their data are again very consistent with our measurements. Roca-Cusachs *et al.* ^18^ used AFM on neutrophils, and found a loss tangent very consistent with our measurements on PLB cells (Fig. 6d).

**Figure 6.**
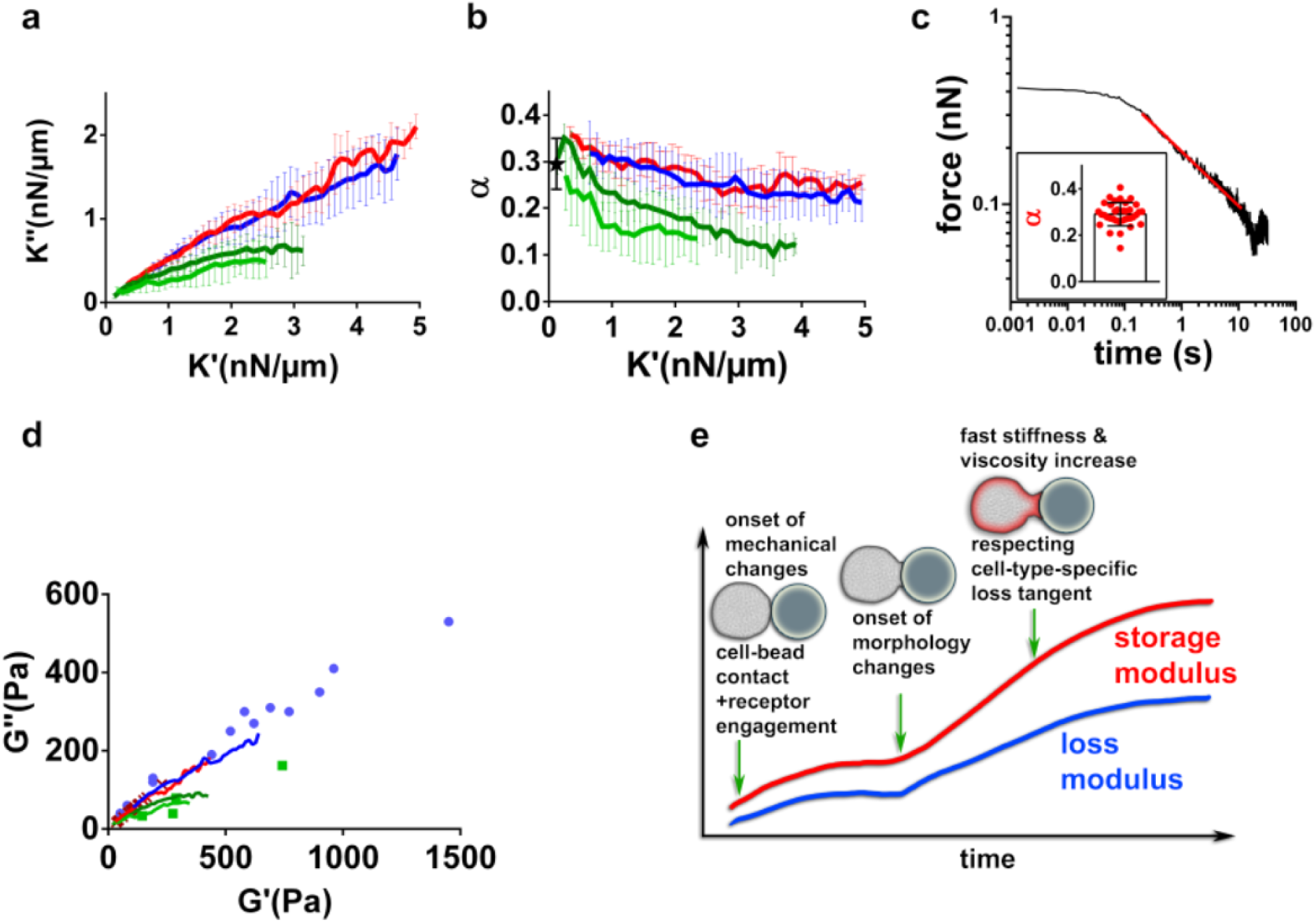
Relationship between elastic and viscous cell properties (**a**) *K” vs. K’* during activation of T cells (red), B cells (blue) and PLB cells (front indentation: dark green, back indentation: light green). Error bar are SDs. (**b**) 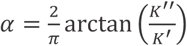 corresponding to the same data as in **a**. The black star is the value of the power-law exponent in force relaxation experiments on resting PLB cells. (**c**) Example of force relaxation curve of a PLB cell. In the log-log plot, a power-law relaxation is identified by the straight line (red), whose slope is the exponent α. Inset: individual α values (cell per dot). (**d**) Loss modulus *G″ vs.* storage modulus *G′* from Bufi *et al.*^62^ (blue dots), Roca-Cusachs *et al.*^18^ (green dots), and Maksym *et al.*^30^(red crosses). Solid lines correspond to data as in **a**. (**e**) Proposed model of mechanical changes during leukocyte activation.

## Discussion

We propose a simple model of leukocyte mechanical changes during activation (fig. 6e): mechanical changes begin seconds after receptor engagement at the leukocyte surface. Cell morphological changes start a few tens of seconds later, concomitant with stiffening and increase in viscosity. Viscous and elastic properties of the leukocyte evolve following a characteristic relationship defined by the loss tangent. The role of these mechanical changes is not clear yet. Cell contractility and cell stiffness are correlated in muscle cells^63–67^; fibroblasts ^33^ and epithelial cancer cells^68^ getting stiffer also get more viscous. Although the inner structure of leukocytes differs from these cell types, the increase in stiffness and viscosity that we observe during activation might be a manifestation of conserved cytoskeletal-based properties among mammalian cells.

Although the molecular details leading to changes in viscoelastic properties are yet to be understood, our observations that cell viscosity increases during leukocyte activation might imply that diffusive properties change during leukocyte activation. The formation of diffusional barriers could be one manifestation of the increase in viscosity that we observe on the whole cell level^69,70^. Diffusion and polarization of organelles such as mitochondria in T cells^71^ might also be impacted by large viscous changes during leukocyte activation. These effects might need to be accounted for in models of leukocyte activation.

Our study highlights that elastic and viscous properties are relevant to understand immune cell mechanics during activation, as well as their relationship in the form of the loss tangent *η*, and how it evolves over time. All the leukocytes that we tested show very large changes in both elastic and viscous properties, but monitoring the time evolution of the loss tangent exhibits a mechanical fingerprint specific to cell types.

It is tempting to speculate that the evolution of the loss tangent during activation can become a valuable tool to discriminate between healthy and pathological leukocytes as already suggested by considerations of static values of *η*^72^. Our study also shows the existence of early mechanical changes, and proposes that information processed at single-receptor scale can be integrated at the cellular scale in only few seconds, thus making mechanical signaling an efficient process worth integrating in mechanistic description of early immunological signaling.

## Materials and Methods

### Micropipette rheometer

The rheometer was build based on the evolution of our microindentation setup^16^ and Micropipette Force Probe^17,37,49^. Micropipettes were prepared as described previously^16,17,37,49^ by pulling borosilicate glass capillaries (Harvard Apparatus, Holliston, MA, USA) with a P-97 micropipette puller (Sutter Instruments, Novato, CA, USA), cutting them with an MF-200 microforge (World Precision Instruments, Sarasota, FL, USA) and bending them at a 45° angle with an MF-900 microforge (Narishige, Tokyo, Japan). Micropipettes were held by micropipette holders (IM-H1, Narishige) placed at a 45° angle relative to a horizontal plane, so that their tips were in the focal plane of an inverted microscope under brightfield or DIC illumination (T cells: TiE, PLB cells: Ti2, Nikon Instruments, Tokyo, Japan) equipped with a 100× oil immersion, 1.3 NA objective (Nikon Instruments) and placed on an air suspension table (Newport). The flexible micropipette was linked to a non-motorized micropositioner (Thorlabs, Newton, NJ, USA) placed on top of a single-axis stage controlled with a piezo actuator (TPZ001; Thorlabs). The bending stiffness k of the flexible micropipette (about 0.2 nN/µm) was measured against a standard microindenter previously calibrated with a commercial force probe (model 406A; Aurora Scientific, Aurora, ON, Canada). Once the activating microbead was aspirated by the flexible micropipette, the cell held by the stiff micropipette was brought in adequate position using a motorized micromanipulator (MP-285; Sutter Instruments). Experiments were performed in glass-bottom Petri dishes (Fluorodish, WPI, Sarasota, FL, USA). Images were acquired using a Flash 4.0 CMOS camera (for T cells), or a SPARK camera (PLB cells), both from Hamamatsu Photonics, Hamamatsu City, Japan. In order to perform rheological experiments, the setup automatically detects the position of the bead at the tip of the force probe at a rate of 400-500 Hz and imposes the position of the base of the flexible micropipette by controlling the position of the piezo stage. The deflection of the force probe is the difference between the position of the bead and the position of the piezo stage. The force applied to the cell is the product of this deflection by the bending stiffness *k*. A retroaction implemented in Matlab (Mathworks) controlling both the camera via the Micromanager software^73^ and the piezo stage moves the latter in reaction to the measurement of the bead position in order to keep a desired deflection of the cantilever: a controlled force is applied to the cell at any given time. Experiments were performed at room temperature to avoid thermal drift. Qualitative behavior of T cell and PLB cells is unchanged at 37°C (not shown). We quantified the fluid drag due to micropipette translation (Suppl. Mat. 11) and the potential effect of fast cell deformation on our rheological measurements (Supp. Mat. 12). *Data analyzis.* Rheological measurements were analyzed by post-processing using a custom-made Python code. During force modulation sinusoidial *x_tip_* signal was fitted every 0.5-1 period interval over a window that was 1.5-2.5 period long (i.e. every 0.5-1 second with a window of 1.5-2.5 seconds for a frequency of force modulation *f*=1 Hz) using a function equal to a linear trend with an added sinusoidal signal (two free parameters for the linear trend, two for the sinusoidal signal when imposing the frequency *f*=1 Hz). Best fit was determined using classical squared error minimization algorithm in Python.

### T cell experiments

#### Cells and reagents

Mononuclear cells were isolated from peripheral blood of healthy donors on a Ficoll density gradient. Buffy coats from healthy donors (both male and female donors) were obtained from Etablissement Français du Sang (Paris, France) in accordance with INSERM ethical guidelines. Human total CD4+ isolation kit (Miltenyi Biotech, 130-096-533) was used for the purification of T cells. Isolated T-cells were suspended in FBS: DMSO (90%:10% vol/vol) and kept frozen in liquid nitrogen. One to seven days before the experiment the cells were thawed. Experiments were performed in RPMI 1640 1x with GlutaMax, supplemented with 10% heat-inactivated fetal bovine serum (FBS) and 1% penicillin-streptomycin (all from Life Technologies ThermoFisher Scientifc, Waltham, MA), and filtered with (0.22 µm diameter-pores, MerckMillipore).

#### Beads

Dynabeads Human T-Activator CD3/CD28 for T Cell Expansion and Activation from Gibco, were purchased from Thermo Fisher Scientific (Carlsbad, CA).

### B cell experiments

#### B cells

Primary B lymphocytes were isolated from spleens of adult C57BL/6J mice (10- to 20-week-old), purified by negative selection using the MACS kit (130-090-862) in the total splenocytes. Animal procedures were approved by the CNB-CSIC Bioethics Committee and conform to institutional, national and EU regulations. Cells were cultured in complete RPMI 1640-GlutaMax-I supplemented with 10% FCS, 1% penicillin-streptomycin, 0.1% b-mercaptoethanol and 2% sodium pyruvate (complete RPMI).

#### Beads

Silica beads (5 × 10^6^; 5 μm diameter; Bangs Laboratories) were washed in distilled water (5,000 rpm, 1 min, RT), and incubated with 20 μl 1,2-dioleoyl-PC (DOPC) liposomes containing GPI-linked mouse ICAM-1 (200 molecules/μm^2^) and biotinylated lipids (1000 molecules/μm^2^) (10 min, RT), and washed twice with beads-buffer (PBS supplemented with 0.5% FCS, 2 mM MgCl2, 0.5 mM CaCl2, and 0.5 g/L D-Glucose). Then, lipid-coated beads were incubated with AF647-streptavidin (Molecular Probes) (20 min, RT) followed by biotinylated rat anti-μB; light chain antibody (clone 187.1, BD Biosciences) (20 min, RT), used as surrogate antigen (su-Ag) to stimulate the B cell receptor. Control beads were coated with ICAM-1-containing lipids only (no su-Ag was added). The number of molecules/μm^2^ of ICAM-1 and su-Ag/biotin-lipids was estimated by immunofluorometric assay using anti-ICAM-1 or anti-rat-IgG antibodies, respectively; the standard values were obtained from microbeads with different calibrated IgG-binding capacities (Bang Laboratories). The lipid-coated silica beads were finally resuspended in complete RPMI.

### PLB cell experiments

#### Cells and reagents

The human acute myeloid leukemia cell line PLB-985 was cultured in RPMI 1640 1X GlutaMax (Gibco) supplemented with 10% heat-inactivated FBS and 1% penicillin/streptomycin, and filtered using a 0.22 µm-diameter pore filter (MerckMillipore). Cells were passaged twice a week and differentiated into a neutrophil-like phenotype by adding 1.25 % (v/v) of dimethyl sulfoxide (DMSO, Sigma-Aldrich) to the cell suspension the first day after passage and, in a second time, three days after while changing the culture media^74^. Before experiments, 2000 U/ml IFN-γ (Immuno Tools) was added into the cell culture flask, cells were centrifuged for 3 min at 300g and resuspended in filtered HEPES medium (140 mM NaCl, 5 mM KCl, 1 mM MgCl2, 2 mM CaCl2, 10 mM HEPES pH 7.4, 1.8 mg/mL glucose, and 1% heat-inactivated FBS). The medium was filtered using a 0.22 µm-diameter pore filter (Merck-Millipore).

#### Beads

20-µm diameter polystyrene microbeads at 10^6^ beads/mL (Sigma-Aldrich) were washed three times by centrifugation at 16000 g for 3min and resuspended in Dulbecco’s Phosphate Buffered Saline (DPBS, Gibco) filtered with 0.22 µm diameter-pores (MerckMillipore). Beads were then incubated overnight at room temperature with 5% (w/v) Bovine Serum Albumin (BSA, Sigma-Aldrich) in DPBS. Beads were washed again three times by centrifugation at 16000 g during 3min, resuspended in DPBS, and incubated with 1:500 anti-BSA rabbit antibody at 3.5 mg/mL (Sigma-Aldrich, ref. B1520) in DPBS for an hour at room temperature. Beads were washed three times by centrifugation at 16000g for 3 min with DPBS and resuspended in DPBS at 10^6^ beads/mL before use.

### Atomic Force Microscope and Single Cell Force Spectroscopy

#### Cells and reagents

3A9m T cells were obtained from D. Vignali^75^ and cultured in RPMI completed with 5 % FBS, 10 mM Hepes, 10mM Sodium Pyruvate in 5 % CO2 atmosphere. COS-7 APCs were generated as previously described ^54^ by stably co-expressing the α and the β chains of the mouse MHC class II I-Ak, cultured in DMEM (5 % FBS, 1mM Sodium Pyruvate, 10 mM Hepes, and geneticin 10µg/ml). Cells were passaged up to three times a week by treating them with either Trypsin/EDTA or PBS 1X (w/o Ca2+/Mg2+), 0.53mM EDTA at 37°C for up to 5 min.

The anti CD45 antibodies used for this study were produced from the hybridoma collection of the CIML (namely H193.16.3)^55^. Briefly, cells were routinely grown in complete culture medium (DMEM, 10% FBS, 1mM sodium pyruvate) prior to switching to the expansion and production phase. Hybridomas were then cultured in DMEM with decreasing concentrations of low immunoglobulin fetal bovine serum down to 0.5%. Cells were then maintained in culture for 5 additional days enabling immunoglobulin secretion prior to supernatant collection and antibody purification according to standard procedures.

The expression of TCR and CD45 molecules on T cells and MHC II molecules on COS-7 APC was assessed once a week on a flow cytometer (anti-TCR PE clone H57.597, anti-CD45 Alexa Fluor 647 clone 30F11 were purchased from BD Pharmingen). The count and the viability, with Trypan blue, were assessed automatically twice by using Luna automated Cell Counter (Biozym scientific). Mycoplasma test was assessed once a month. Culture media and PBS were purchased from Gibco (Life Technologies). PP2 (Lck inhibitor) was purchased from Calbiochem.

#### T Cell immobilization

Culture treated, sterile, glass bottom Petri dishes (World Precision Instruments Fluorodish FD35-100) were incubated with 50µg/ml anti-CD45 antibody for 1 hour at room temperature. The surfaces were extensively washed with PBS 1X prior to a last wash with HBSS 1X 10 mM Hepes. The surfaces were kept wet with HBSS 1X 10 mM Hepes before seeding T cells.

#### T cell preparation

T cells were counted, spinned and resuspended in HBSS 1X 10 mM Hepes and were kept for one hour at 37 °C 5% CO2 for recovery prior to seeding. The cells were seeded at room temperature and after 30 minutes a gentle wash was done by using two 1mL micropipettes in order to maintain, as much as possible, a constant volume in the Petri dish and hence avoid perturbations by an important flow. After one hour incubation the Petri dish was mounted in a Petri dish Heater system (JPK Instruments) to maintain the temperature of the sample at ~ 37°C. 10 minutes before experiments, 10µM (final concentration) of Lck kinase inhibitor, PP2 drug was added and homogenized in the sample. We kept this drug present all along the experiments.

#### COS-7 APC preparation

The day before the experiment, cells were incubated with the peptide of interest with a final concentration of 10µM allowing 100% occupation of MHC II molecules. Before the experiment the cells were detached from the cell culture plates by removing the cell medium, washed once with PBS 1X and then by a 0.53 mM EDTA treatment for 5 minutes at 37°C 10% CO2. The cells were resuspended in HBSS 1X 10mM Hepes and were allowed to recover before to be seeded into the sample. The peptide p46.61 (which is CD4 dependent) was purchased from Genosphere with a purity >95%.

#### AFM setup

The setup has been described in great detail elsewhere^76^. Measurements were conducted with an AFM (Nanowizard I, JPK Instruments, Berlin) mounted on an inverted microscope (Zeiss Axiovert 200). The AFM head is equipped with a 15 μm *z*-range linearized piezoelectric scanner and an infrared laser. The set-up was used in closed loop, constant height feedback mode^53^. Bruker MLCT gold-less cantilevers (MLCT-UC or MLCT-Bio DC) were used in this study. The sensitivity of the optical lever system was calibrated on the glass substrate and the cantilever spring constant by using the thermal noise method. Spring constant were determined using JPK SPM software routines *in situ* at the start of each experiment. The calibration procedure for each cantilever was repeated three times to rule out possible errors. Spring constants were found to be consistently close to the manufacturer’s nominal values and the calibration was stable over the experiment duration. The inverted microscope was equipped with 10x, 20xNA0.8 and 40xNA0.75 lenses and a CoolSnap HQ2 camera (Photometrics). Bright field images were used to select cells and monitor their morphology during force measurements through either Zen software (Zeiss) or Micro-Manager^73^.

#### APC/T cell force measurements

*Lever decoration.* To make the cantilevers strongly adhesive, we used a modified version of our previous protocols^77^. Briefly, cantilevers were activated using a 10min residual air plasma exposure then dipped in a solution of 0.25mg/mL of Wheat Germ Agglutinin (WGA) or 0.5 mg/mL of Concanavalin A in PBS 1x for at least one hour. They were extensively rinsed by shaking in 0.2µm filtered PBS 1x, stored in PBS 1x at 4°C. Before being mounted on the glass block, they were dipped in MilliQ-H_2_O in order to avoid the formation of salt crystals in case of drying and hence alteration of the reflection of the laser signal. *APC capture.* In separate experiments, we used side view and micropipette techniques, and we observed that (i) this allows the presented cell to be larger than the lever tip, excluding any unwanted contact of T cell with WGA and (ii) the binding is resistant enough to ensure that any rupture event recorded is coming from the cell/cell interface (Suppl. Fig. S6). After calibration, a lectin-decorated lever is pressed on a given COS7 cell for 20 to 60 seconds with a moderate force (typically 1-2nN), under continuous transmission observation. Then, the lever is retracted far from the surface and the cell is let to recover and adhere/spread for at least 5 minutes before starting the experiments (Suppl. Fig. S6). *SCFS experiment.* The surrogate APC is then positioned over a desired T cell. Because the lever is not coated with gold, one can finely position the probe over the target. Force curves are then recorded using the following parameters: max contact force 1nN, speeds 2µm/sec, acquisition frequency 2048 Hz, curve length 10µm and contact time 60 sec. At the end of the retraction, lateral motion of the cells is made by hand until no more force is recorded by the lever, signaling that full separation was achieved (Suppl. Fig. S6). One APC cell was used to obtain one force curve on at least three different T cells, and 3 APCs at least were used for each condition. No apparent bias was detected in the data when carefully observing the succession of the measures for a given APC, for a given lever and for a set of T cells. *Data processing & Statistics.* Force curves were analyzed using JPK Data Processing software on a Linux machine. Each curve was processed manually, except for calculation and plotting of mean±SD for relaxation curves where *ad-hoc* python scripts were used. Data were processed using Graphpad Prism (v7) on a Windows 7 machine. On graphs, data are presented as scatter dot plots with median and interquartile range. A data point corresponds to a force curve obtained for a given couple T cell/APC, except for the small detachment events where one data point corresponds to one such event (all events were pooled). Significance was assessed using Mann-Whitney tests in GraphPad Prism v7 with p<0.05 *, <0.01 **, <0.001 ***, and **** below; not significant otherwise.

## Acknowledgements

This work has benefited from the financial support of the LabeX LaSIPS (ANR-10-LABX-0040-LaSIPS) managed by the French National Research Agency under the "Investissements d'avenir" program (n°ANR-11-IDEX-0003-02), from a CNRS PEPS grant, from Ecole Polytechnique, and from an endowment in cardiovascular bioengineering from the AXA Research Fund. Part of this work was supported by funding from Prise de Risques CNRS (PHP), the ANR JCJC « DissecTion » (PHP), and from the Labex IN-FORM (ANR-11-LABX-0054) and A*MIDEX project (ANR-11-IDEX-0001-02) funded by the “In-vestissements d’Avenir” French Government program managed by the French National Research Agency (ANR) (to Inserm U1067 Lab & CIML, and as PhD grant to A.S.). Part of this work was also supported by institutional grants from INSERM, CNRS and Aix-Marseille University to the CIML. *Material or technical help*: R. Balland (Sensome) for RBCs; M. Pélicot-Biarnes (LAI); L. Borge (PCC cell culture facility); S. Mailfert / M. Fallet / PICSL imaging facility of the CIML (ImagImm), member of the national infrastructure France-BioImaging supported by the French National Research Agency (ANR-10-INBS-04); A. Formisano (He/Marguet lab). *Discussion and support*: D. Gonzalez-Rodriguez (LCP-A2MC, Univ. De Lorraine), R Roncagalli (CIML); P. Recouvreux (IBDML); L. Limozin, F. Rico (LAI). Companies : F. Eghiaian, T. Plake & JPK Instruments (Berlin, Germany), now part of Bruker, for continuous support and generous help ; Zeiss France for support. SVMC and YRC have benefited from the financial support of an FPI contract (BES-2014-068006) and the grant BFU2013-48828-P from the Spanish Ministry of Economy (MINECO).

## Author contributions

P.-H.P., Y.H. and J.H. designed experiments; A.Z., S.R.M.C., A.S., P.-H.P. and J.H. performed experiments; A.Z., A.S., P.-H.P. and J.H. analyzed the data; A.B., S.D., E.H., S.D.-C., H.-T.H., A.I.B., Y.R.C., Y.H., C.H. and O.N. contributed material; P.-H.P. and J.H. wrote the manuscript with helpful critical reading by the other authors.

## Competing financial interests

The authors declare no competing financial interests.

