## Supplementary Material for "Single-cell immuno-mechanics: rapid viscoelastic changes are a hallmark of early leukocyte activation"

[1] Laboratoire d'Hydrodynamique (LadHyX), Ecole polytechnique, CNRS UMR7646, Institut Polytechnique de Paris, route de Saclay, 91128 Palaiseau Cedex, France.
[2] Laboratoire de Chimie Physique, CNRS, Université Paris-Sud, Université Paris-Saclay, 91405, Orsay, France.
[3] B cell Dynamics Laboratory, Department on Immunology and Oncology, Centro Nacional de Biotecnología (CNB)-CSIC, Madrid, Spain.
[4] Aix Marseille University, LAI UM 61, Marseille, F-13288, France.
[5] Inserm, UMR_S 1067, Marseille, F-13288, France.
[6] CNRS, UMR 7333, Marseille, F-13288, France.
[7] Institut Curie, PSL Research University, INSERM U932, Integrative analysis of T cell activation team, 26 rue d'Ulm, 75248, Paris Cedex 05, France.
[8] Aix Marseille University, CNRS, INSERM, CIML, Marseille, France.

### Supplementary Material

#### Supplementary movies

Movie S1 | Activation of three types of leukocytes studied with the micropipette rheometer (top: T cell, middle: B cell, bottom: PLB cells). All bars represent 5 µm. Time is in minutes:seconds.

Movie S2 | Validation of the micropipette rheometer by performing microindentation measurements on a red blood cell. The bar represents 5 µm. Time is in minutes:seconds.

Movie S3 | Modified setup to indent the "back" of a PLB cell while the cell phagocytoses an activating bead on its "front". The bar represents 10 µm. Time is in minutes:seconds.

Movie S4 | cycles PLB (cycles_8um_181207_cell8 ou cell2). Cyclic indentation experiments to measure directly the Young's modulus over time of a PLB cell phagocytoses an activating bead. The bar represents 10 µm. Time is in minutes:seconds.

Movie S5. Modified setup using an auxiliary pipette to bring an activating microbead in contact with a T cell on its "side". The bar represents 5 µm. Time is in minutes:seconds.

#### Supplementary material 1 | Measuring initial cell Young's modulus.

The Hertz model[1] leads to the following relationship between the applied compressive force and cell indentation δ : $F = \frac{4}{3}\frac{E_{Young}}{1-\nu^2}\sqrt{R_{eff}}\delta^{\frac{3}{2}}$, where $E_{Young}$ is the effective Young's modulus of the cell, $R_{eff}$ is an effective radius given by $1/R_{eff} = 1/R_{cell} + 1/R_{bead}$, with $R_{cell}$ the cells radius and $R_{bead}$ the bead radius; ν is the Poisson ratio of the cell taken as 0.5, i.e. the cell is considered incompressible [2,3].

#### Supplementary material 2 | Complex stiffness.

We apply a force $F(t) = \langle F\rangle + \Delta F\cos(\omega t)$ to the leukocyte (angular frequency $\omega = 2\pi f$, frequency $f$ = 1 Hz) and we measure a resulting oscillatory change in position of the tip of the flexible micropipette

$x_{tip}(t) = \langle x_{tip} \rangle + \Delta x_{tip} \cos(\omega t - \varphi)$, of average value $\langle x_{tip} \rangle$, amplitude $\Delta x_{tip}$, and phase lag $\varphi$. We consider a complex formalism where the force writes as $F^*(t) = \langle F \rangle + \Delta F e^{i\omega t}$ and the tip position writes as follows: $x^*_{tip}(t) = \langle x_{tip} \rangle + \Delta x_{tip} e^{i(\omega t + \varphi)}$. We then define the complex stiffness $K^*$ by writing $F^*(t) - \langle F \rangle = K^*\left(x^*_{tip}(t) - \langle x_{tip} \rangle\right)$, this leads to $\Delta F e^{i\omega t} = K^* \Delta x_{tip} e^{i(\omega t - \varphi)}$ where we can divide both terms of the equality by the complex exponential term $e^{i\omega t}$ to get $\Delta F = K^* \Delta x_{tip} e^{-i\varphi}$. Now by multiplying both terms of the equality by $e^{i\varphi}$, decomposing terms in their real and imaginary parts ($K^* = K' + iK''$ and $e^{i\varphi} = \cos\varphi + i\sin\varphi$), and equating real and imaginary parts we obtain two relationships:
$\Delta F \cos\varphi = K' \Delta x_{tip}$ and $\Delta F \sin(\varphi) = K' \Delta x_{tip}$ finally leading to :

$$K' = \frac{\Delta F}{\Delta x_{tip}} \cos\varphi$$

$$K'' = \frac{\Delta F}{\Delta x_{tip}} \sin\varphi.$$

#### Supplementary material 3 | Validation with red blood cells.

We measured large viscous components in leukocytes (*K''* lower but comparable to *K'*, and large equivalent viscosities, fig. 2). These were measured based on the phase lag $\varphi$ between the applied force and the resulting deformation of the cell. In order to validate our technique against a consensually well-characterized cell type, we applied it to red blood cells (RBCs). There are several models predicting the stiffness of a RBC (quantified by *K'*) held aspirated by a micropipette and used as a spring in a Biomembrane Force Probe (BFP)[4–6]; these models were recently compared by Ju and Zhu[7]. Human RBCs (or pig RBCs as tested here) are devoid of nucleus and of the complex and space-occupying organelles present in leukocytes, so we expected the viscous part of RBC complex stiffness to be comparatively much smaller than in a leukocyte. We quantified both *K'* and *K''* in RBCs aspirated in a micropipette with a controlled aspiration pressure (Suppl. Fig. S1, movie S1). In the BFP, a bead is stuck to the tip of the red blood cell to build the force probe. The corresponding model requires measurement of the geometry of the RBC (outer diameter, diameter of the part inside the micropipette), and RBC-bead contact radius. In our case the indenting bead played the role of the BFP bead. As the RBC-bead contact radius is difficult to measure accurately with light microscopy, we took the value given by the Hertz model knowing the applied compressive force and the shape of the indenter and the RBC under zero force. We obtained a good agreement between the measured *K'* and the predicted *K'* (within ~20%, Suppl. Fig. S1b). Values for *K''* were very low, and did not increase with increasing *K'* as opposed to what we saw in leukocytes (Suppl. Fig. S1c). These experiments confirm that the large viscous components of leukocyte complex stiffness are due to actual viscous behavior, which as expected is absent when measuring RBCs.

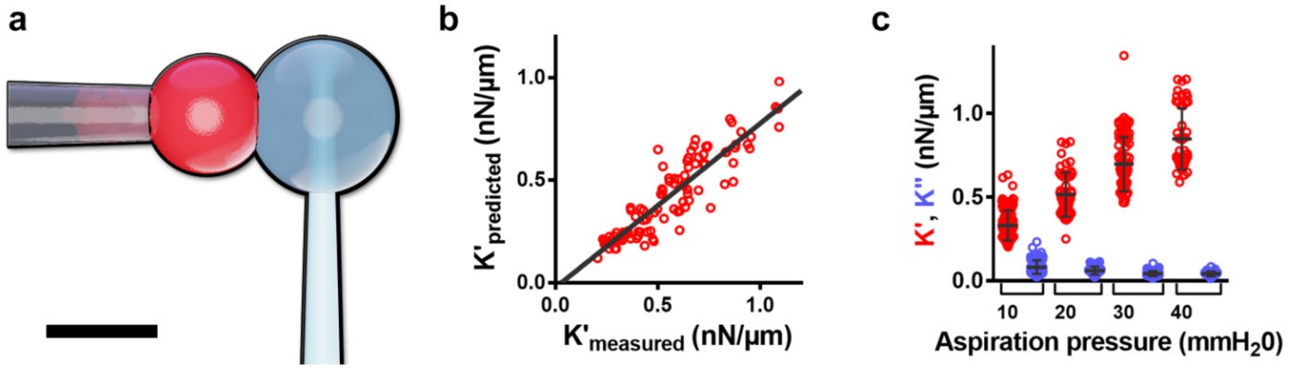


**Supplementary Figure S1 |** Viscoelastic properties of red blood cells. (**a**) Schematic view of the experiment. A red blood cell (RBC) is held by a rigid micropipette (left) under a controlled aspiration pressure. A glass microindenter is used to apply an oscillatory force to the RBC. The bar represents 5 µm. (**b**) Predicted *K'* vs measured *K'*. A linear fit of the data leads to predicted $K'_{predicted} = 0.80 K'_{measured}$ - 0.02 nN/µm (n=3 experiments, N=18 cells, each dot is the mean of 3 oscillation periods repeated over the same cell). (**c**) Elastic (*K'*) and viscous (*K''*) part of aspirated RBCs as a function on the aspiration pressure applied in the holding micropipette. Error bars are standard deviations (SD).

#### Supplementary material 4 | Measuring initial storage modulus *E'* and loss modulus *E''*.

From the initial compression of the cell by the bead, when the leukocyte is still round, we use the Hertz model (see methods in main text) to measure the Young's modulus of the cell. To extract *E'* and *E''* from the first seconds of the application of force modulation, we use a linearization of the Hertz model as done elsewhere[8–13] leading to:

$E' \sim \frac{1-\nu^2}{2}\frac{K'}{\sqrt{R_{eff}\langle x_{tip}\rangle}}$ and $E'' \sim \frac{1-\nu^2}{2}\frac{K''}{\sqrt{R_{eff}\langle x_{tip}\rangle}}$, where $K' = \frac{\Delta F}{\Delta\delta}\cos\varphi$ and $K'' = \frac{\Delta F}{\Delta\delta}\sin\varphi$, where ΔF is the force amplitude, $\Delta x_{tip}$ the resulting oscillation amplitude, and φ the phase lag., $\nu$ the Poisson ratio of the cell taken as 0.5, and $R_{eff}$ the effective radius of the indenting bead-cell system (Suppl. Mat. 1).

#### Supplementary material 5 | Measuring an effective Young's modulus during activation.

Measuring the equivalent Young's modulus of the cells over time during activation requires to model the geometry of the cells in a way that is adapted to each cell type. We did so for T cells and B cells, but for PLB cells we could directly measure a Young's modulus by using the modified setup described in figure 3a in main text. Here we explain this approach in more detail.

*Deducing an equivalent Young's modulus from E'.* We compared $E_{Young}$ and *E'* measured at an initial time point for T cells, B cells, and PLB cells. *E'* is correlated with, but 1-2 fold larger than $E_{Young}$; we introduced an ad hoc cell type-dependent constant *C* such that $E' = CE_{Young}$ ; *C* is 1.6±0.1 for CD4, and 1.8±0.1 for B cells. For PLB cells, we used the alternative setup to indent the phagocyte on its "back" in a cyclic way (see figure 4a and main text), which allowed us to measure directly a Young's modulus over time every ~30 seconds (dotted curve in fig. 2l). This leads to a coefficient C = 1.12±0.02 for PLB cells valid even during activation. In the case of T and B cells, we assumed that similar to the case for PLB cells, we could consider that the factor C is rather constant over time. Using this approach, we measured an effective Young's modulus of the three cell types over time (figure 2j-l).

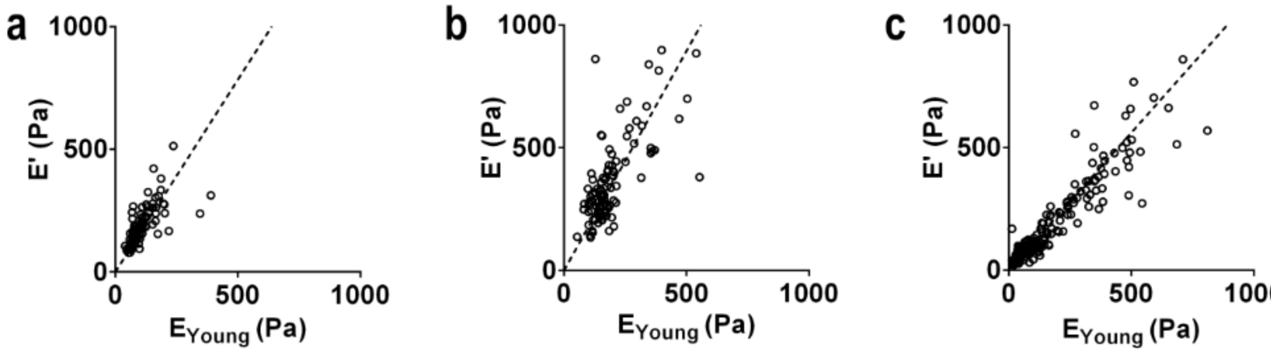


**Supplemental figure S2 |** The equivalent modulus *E'* is proportional to the Young's modulus $E_{Young}$: $E' = CE_{Young}$, where *C* = 1.6±0.1 for T cells (**a**), 1.8±0.1 for B cells (**b**), and 1.12±0.02 for PLB cells (**c**).

**T cells.** During first minutes of activation, T cells develop a cylindrical protrusion[14]. We model the whole cell as a cylinder, which leads to $E_{Young}^{eq} = \frac{1}{C}K'\frac{L}{S}$, where *L* is the length of the cell, *S* the section of the cylinder, and *C*=1.6 the correction factor described above to convert E' in $E_{Young}^{eq}$. The resulting equivalent Young's modulus (fig. 2j) increases until reaching a maximal value that is 4.7±2.6 times its initial value, consistent with the observed increase in *K'* (fig. 2d).

**B cells.** During activation, B cells remain more round than T cells, but still change in dimension and losing the initial cell length makes the Hertz model difficult to use. We thus considered two different estimates for the equivalent Young's modulus $E^{eq}_{Young}$.

**PLB cells.** Activating PLB cells spread on their target (Fig. 2c), requiring yet another model. In order to be able to apply the Hertz model, we used a modified setup where a non-adherent glass bead indents the cell on its "back" (figure 4a) while the cell phagocytoses the microbead on its "front". The measured *K'* and *K''* led to the same $K'_{max}/K'_{init}$ and $K''_{max}/K''_{init}$ ratios as those obtained with front indentation (figure 4b). To compare the variation in K' with the variation in cell tension reported by others[15–18], we show that an increase in tension reflects an increase in K' (Suppl. Mat. 8).

*First approximation.* Starting from the storage modulus $E' \sim \frac{1-\nu^2}{2}\frac{K'}{\sqrt{R_{eff}\langle x_{tip}\rangle}}$, we considered the relationship given by the Hertz model between the average force <F> and average indentation <δ> : $\langle F\rangle = \frac{4}{3}\frac{E_{Young}}{1-\nu^2}\sqrt{R_{eff}}\langle\delta\rangle^{\frac{3}{2}}$, and substituted the resulting expression for <δ>, obtaining:

$E^{eq}_{Young} = (1-\nu^2)(\frac{K'}{C})^{3/2}\left(\frac{1}{6R_{eff}\langle F\rangle}\right)^{1/2}$,with the correction factor C (C=1.8 for B cells) as described in supplementary material 5. The deduced $E^{eq}_{Young}$ at the contact time was much higher than the Young's modulus measured using constant-speed indentation. A careful data inspection showed that after the maximal force of 120 pN was reach during initial compression, when the force switched to the force modulation mode with a compressive average force of 60 pN, the indentation did not decrease as much as predicted by the Hertz model. This can be due to several effects, including adhesion[19], complex visco-elasto-plasticity [20] or poroelasticity [3].

*Second approximation.* In a second approximation we considered that the indentation measured over the first few periods of application of the oscillatory force was constant over time $E^{eq}_{Young} \sim \frac{1}{C}\frac{1-\nu^2}{2}\frac{K'}{\sqrt{R_{eff}\langle x_{tip}\rangle}}$, where $\langle x_{tip}\rangle$ was maintained constant and equal to its initial value. This approximation is an underestimation of $E^{eq}_{Young}$ because $\langle x_{tip}\rangle$ under a given force decreases when cell stiffness increases. This estimation matched the initial value of the Young's modulus, and led to a time evolution comparable to values obtained in T cells (Figure 2).

#### Supplementary material 6 | Spatial localization of viscoelastic changes.

We asked if mechanical changes in T cells during activation depended on the distance to the cell-bead contact area. We used a modified setup, where an auxiliary pipette brings the activating microbead in contact with the cell on its "side": at the equatorial plane (distance between the activating and indenting beads smaller than 5 μm), or close to the tip of the holding pipette, at the farthest possible location from the activating bead (distance between the activating and indenting beads larger than 5μm, Suppl. Fig. S3, Suppl. Movie S5). We computed an equivalent Young's modulus $E^{eq}_{Young}$ as described in supplementary material 5, which led to a good agreement with $E_{Young}$ measured at the initial time. T cells exhibited a different behavior depending on the location of the activating microbead: the closer the indenter was to the activating bead, the larger the maximum value of $E^{eq}_{Young}$ measured during activation. Cell stiffening thus occurs with a different amplitude depending on the location on the cell relative to the contact zone with an activating surface.

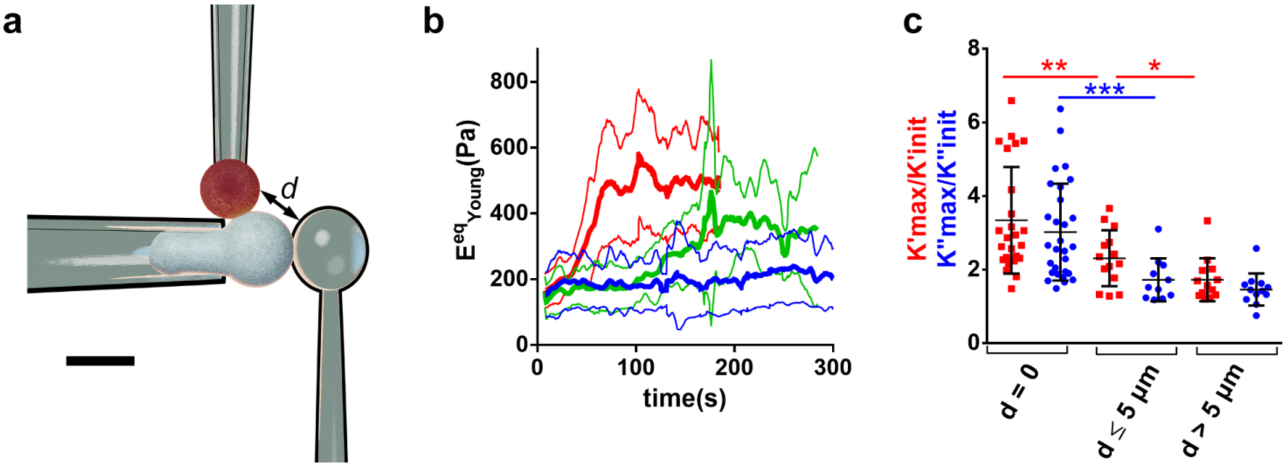


**Supplementary figure S3**| (**a**) Modified setup where an auxiliary (stiff) pipette holds the activating microbead (top, red) to bring it in contact with the T cell at a chosen distance from a microindenter (right) which indents the cell on its "side" while the T cell gets activated. Scale bar is 5 µm. (**b**) Equivalent Young's modulus for three different ranges of distance *d* between the activating bead and the microindenter (red: *d*=0; green: 0 < *d* ≤ 5 µm; blue: *d* > 5 µm). (**c**) $K'_{max}/K'_{init}$ (red) and $K''_{max}/K''_{init}$ (blue) ratios for the same ranges of distance *d* as in panel **b**.

#### Supplementary material 7 | Cell stiffness increases before membrane surface stores are exhausted.

In T cells, at the onset of the faster increase in *K'*, the apparent cell surface area as measured optically by transmitted light optical microscopy has only expanded by 5% relative to its initial value. Cell surface area expands further, up to 11% (fig. 5d) when the cell protrusion is at its largest (Suppl. Fig. S4). This increase of 11% is smaller than the 13% that we previously reported as a "slack" or reserve after which cell effective stiffness starts to increase due to progressive exhaustion of surface membrane stores[21]. We conclude that the increase in *K'* is not directly linked to a limited amount of membrane surface reservoirs, at least not in the early moments of activation.

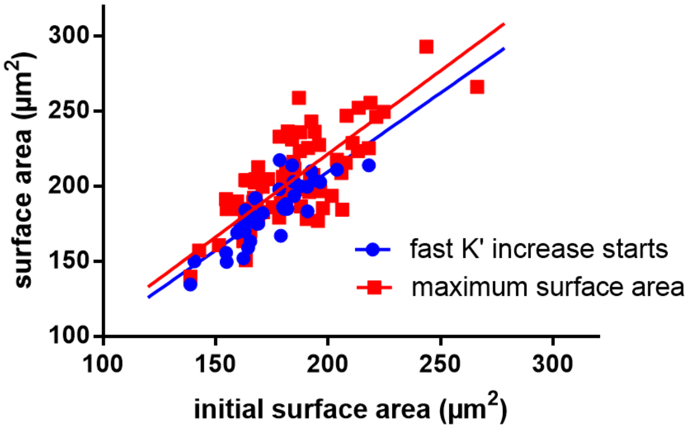


**Supplementary figure S4** | Total apparent cell surface area at two different moments in T cells. In blue: apparent surface area when *K'* starts increasing faster *vs* initial apparent cell surface area of the cell. The straight blue line is a linear fit of slope 1.05± 0.01. In red: maximal apparent surface area *vs* initial apparent cell surface area of the cell. . The straight blue line is a linear fit of slope 1.11± 0.01.

#### Supplementary material 8 | Increase in stiffness corresponds to an increase in cell tension.

The moment at which *K'* starts increasing more rapidly corresponds to the moment at which the tail of the cell in the holding micropipette starts retracting (Suppl. Fig. S5), which is a consequence of an increase in cell tension. Indeed, when modelling the cell as having a cortical tension (as described in the liquid core-cortical shell model of leukocytes[22]), the moment at which the recoil happens is the moment when cell tension $T$ exceeds the level that can be equilibrated by the aspiration pressure inside the micropipette holding the cell. The so-called critical pressure relative to the exterior of the micropipette that is needed to aspirate the tail in the micropipette up to a length equal to the pipette diameter is given by the Laplace law [22]: $\Delta P_c = 2T\left(\frac{1}{R_{pipette}} - \frac{1}{R_{cell}}\right)$. This is an underestimation of the cell tension (we will use here the term cell tension, which represents a combination of membrane tension and cortical tension[23]) that keeps increasing during the recoil process. Cell stiffness K' and cell tension are not independent. The balance between the external pressure and the pressure applied by the indenter on the side, and the cell internal pressure and its tension on the other side (similar to the calculation by Rosenbluth *et al.*[24]) can lead to $K' = \pi\gamma$ as shown by Cartagena-Rivera *et al.* [25] At the moment at which the cell starts its retraction from the holding pipette, we measure $K'$ = 0.9±0.4 nN/µm for T cells. This provides a first estimation of cell tension $T$=$K'/\pi$=0.28 nN/µm. On the other hand, Laplace law applied at the moment the cell starts its retraction leads to $P_{cell} = P_{ext} + 2T\left(\frac{1}{R_{pipette}} - \frac{1}{R_{cell}}\right)$, so that numerically $T \sim \frac{1}{2}\frac{60Pa}{(0.6-0.3)\mu m^{-1}} \sim$ 0.3 mN/m, in excellent agreement with the previous estimate of cell tension[15-18], confirming that K' is a reliable indicator of cell tension.

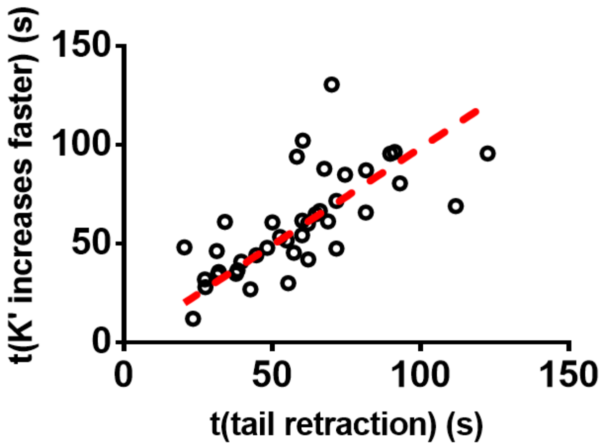


**Supplementary figure S5|** Moment at which *K'* starts increasing faster *vs* time at which the T cell tail in the holding micropipette starts retracting. The dashed line is a linear fit of slope 0.99±0.05.

#### Supplementary material 9 | Cell-cell contact mechanics measured with AFM

We immobilized 3A9 murine CD4+ T cell hybridomas on an aCD45-coated surface on which density does not prevent efficient activation of the T cells by soluble aCD3 or peptide loaded APCs in suspension (not shown). To prevent the rise of signaling cascades, our measurements were performed in the presence of the pan-Src kinase inhibitor PP2. In addition, our surrogate APCs have been extensively documented to be devoid of co-stimulation and adhesion molecules that could interfere with TCR/pMHC interactions[26-28].

The contact mechanics at initial time was, as expected, not modified by the presence or nor absence of peptide on the APC: the slopes of force vs. tip sample separation are essentially the same (Suppl. Fig. S6d).

Then, by measuring the forces during the separation of APC and T cell, we observed that the maximal detachment force (Suppl. Fig. S6e), in parallel with the observed number (Suppl. Fig. S6f) of the small detachment events was significantly increased when a peptide was present, with an effect shown to be peptide dependent, but the magnitude of these events (Suppl. Fig. S6f) was essentially not different, in agreement with what was observed before in AFM single molecule experiments[29]. The forces for the small events, potentially single pMHC/TCR detachments, were on the same range as the one reported earlier. The force we called "offset" (Suppl. Fig. S6c) that we observed due to our limited pulling range, was also peptide dependent (Suppl. Fig. S6f), indicating that non-detached molecules were present and that their number was influenced by the peptide, which has to be paralleled with the previous observation on the number of small events, therefore being an underestimate of the number of molecular bridges to be separated upon cell/cell detachment. Of note, we never observed detachment occurring at the interfaces between aCD45/T cell and/or lectin/APC, the cells being essentially immobile, even when we used large relative, lateral displacement to achieve full separation at the end of the SCFS cycle. All of these elements support the peptide specificity of our SCFS experiments.

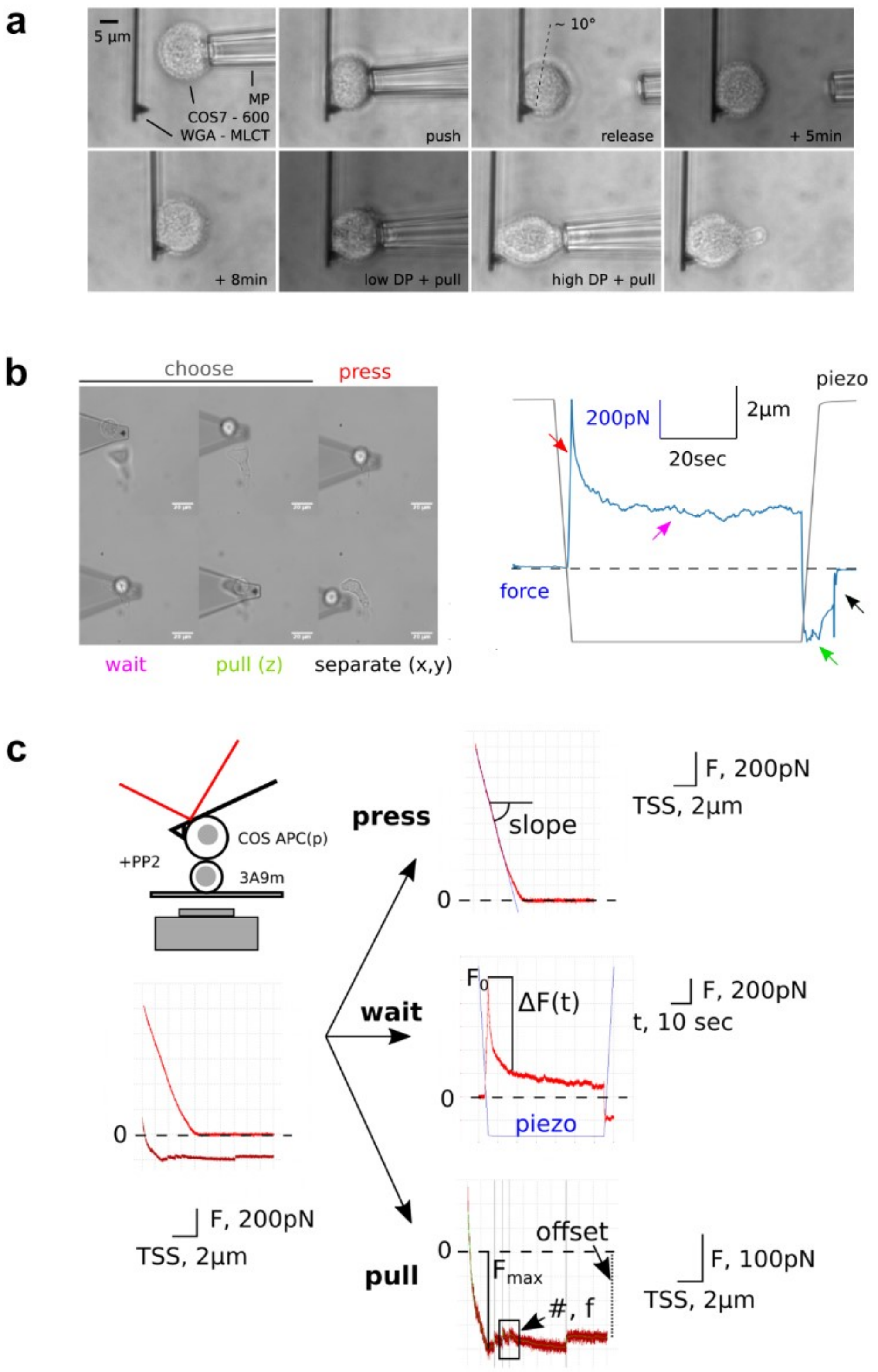

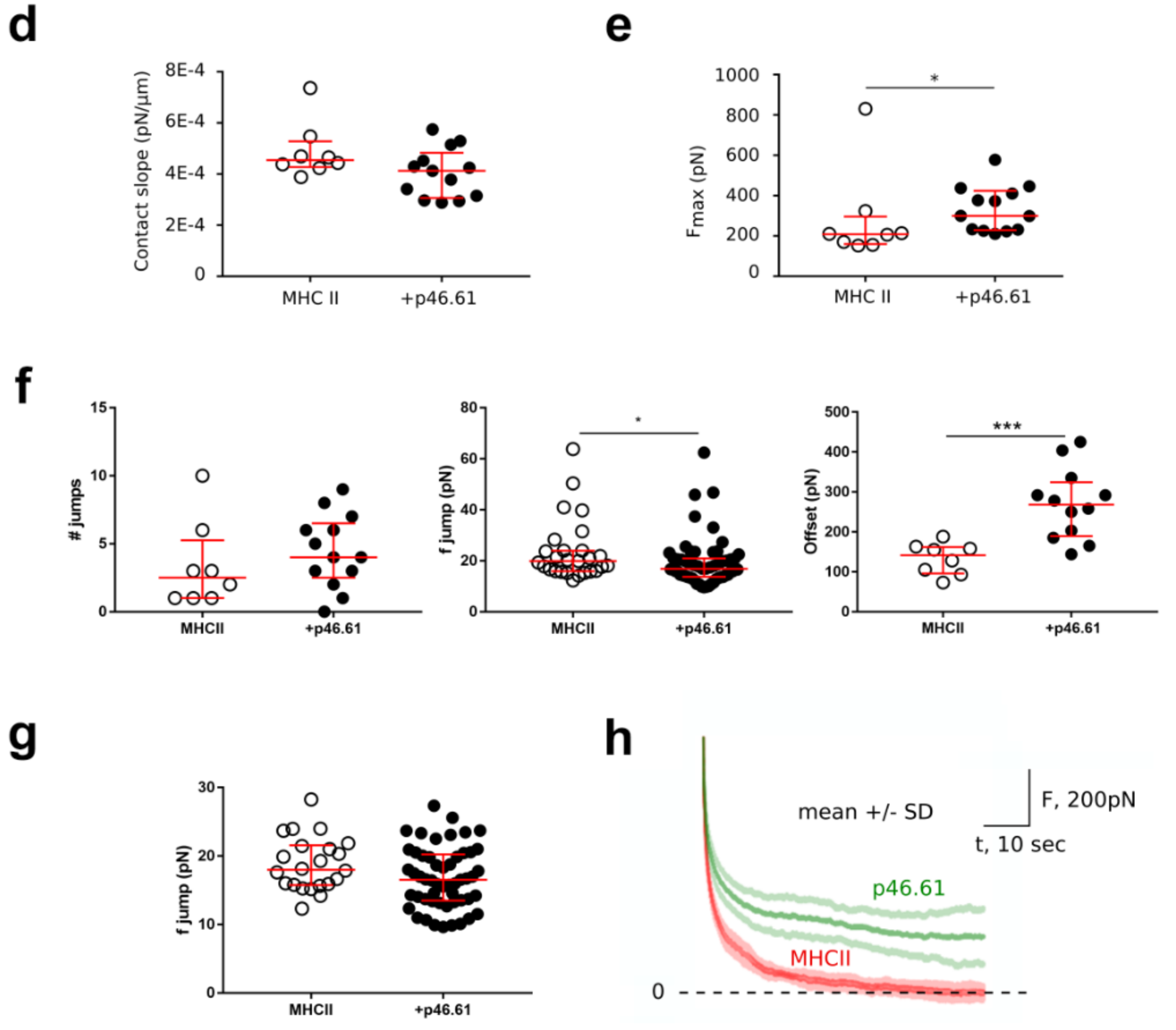


**Supplementary figure S6 |** (**a**) Lateral view of a sequence of micro-mechanical tests of APC attachment to a lectin coated lever. Gentle contact of a micropipette carried APC is followed by an adhesion time (typically several minutes). The adhesion is then tested by strongly aspirating the APC in the micropipette and pulling by moving the pipette away. We never recorded any large variation of the contact zone or separation of the cell from the lever, even after huge aspiration, motion and deformation (see the tongue, which then is "re-adsorbed" by the APC over time). Note that the side observation allows to assess that the tip of the lever, being coated with lectin, never protrudes from the APC, hence is never in contact with the T cell (since, of note, lectins may interfere with T cell activation). Bar = 5 µm. (**b**) Sequence of SCFS, after the APC being adhered to the lever. Left, micrographs; right, force vs. time (blue) and piezo vs. time (grey). After positioning the APC over a T cell, the two are brought into contact, let interact for 60 sec, then pulled separated for the limited piezo range of the system. Full separation is obtained by lateral relative displacement of the two cells, until no more force is recorded by the lever deflection, indicating also that the lever drift was minimal if not null. The color coding of the arrows on the plot indicate the moment. Bar = 10µm. (**c**) APC/T cell Single Cell Force Spectroscopy (SCFS) experiments. Cells are brought into contact under a prescribed force of 1nN, then the AFM piezo keeps its position constant for a duration of 60 sec before the cells are pulled apart. (**d-g**). Effect of the presence of the peptide on the APC. (**d**) Slope (of force *vs.* tip-sample separation (TSS) when putting the cells into contact, as a marker of cell/cell mechanics. No significant difference is recorded ; $p$=0.089. (**e**) Maximal separation force ($p$=0.0246). (**f**) Number of small detaching events, as seen as force jumps, per curve ; even if the distributions are not significantly different (p=0.2884), the case where peptide is implicated presents higher number of small events.. Small jump force magnitude, each of the recorded event being pooled for a given condition, regardless of its nature (adhesion or tether pulling,[30]). Weak variation ($p$=0.0243) and amplitude of these forces as a function of the presence of the peptide in the MHC is consistent with previous single molecule data on T cell murine hybridomas[29]. Offset of the force to the baseline at the end of the accessible pulling distance, which may encompass several non-broken bonds / membrane tethers[30]

($p$=0.005). (**g**) Distribution of force jumps that are below 30pN, which are more likely to be only single molecule separation, potentially excluding tethers and/or multiple events, in regard to Puech *et al.*[29]. No significant difference is then recorded ($p$=0.0631). (**h**) Mean ± SD representation of data in figure 5b in main text.

#### Supplementary material 10 | Structural damping

The loss tangent is a concept employed in the structural damping model that has been successfully applied to describe cell mechanical properties[31]. It relies on the fact that, similar to what is observed in soft glassy materials[32,33], the cell's stress relaxation is a power-law such that $\sigma(t) = \mu\delta(t) + \sigma_0 \left({}^{t}/_{t_0}\right)^{1-x}$ where $\sigma_0$ is the stress per unit strain, $\mu$ a Newtonian viscous term, and $\delta$ the Dirac delta function. The Fourier transform of this temporal stress relaxation leads to the complex modulus $G^*(\omega) = G'(\omega) + iG''(\omega) = G_0 \left(\frac{\omega}{\Phi_0}\right)^{x-1} (1 + i\bar{\eta})\Gamma(2 - x) \cos\frac{\pi}{2}(x - 1) + i\omega\mu$, where $G_0$ and $\Phi_0$ are respectively scale factors for stiffness and frequency, $\Gamma$ is the gamma function, $\mu$ is a Newtonian viscosity, $i$ the unit imaginary number $\sqrt{-1}$, $\bar{\eta}$ is the so-called hysteresivity or the structural damping coefficient [31,3435]. There is now an abundant literature on observed power-law stress relaxation functions in several cell types [9,34–39]. From this last equation we can write the loss tangent $\eta = \frac{E''}{E'} = \frac{E_0\left(\frac{\omega}{\Phi_0}\right)^{\alpha}\bar{\eta}\Gamma(1-\alpha)\cos\frac{\pi}{2}\alpha + \omega\mu}{E_0\left(\frac{\omega}{\Phi_0}\right)^{\alpha}\Gamma(1-\alpha)\cos\frac{\pi}{2}\alpha}$, where $\bar{\eta} = \tan\left(\frac{\pi}{2}\alpha\right)$, and if we consider as Fabry *et al.*[31] that the Newtonian term can be neglected at low frequency (for frequencies of a few Hertz and less – we use 1 Hz-frequency), $\eta$ tends to $\bar{\eta}$ at low frequency. Following this model, the loss tangent $\eta$ is thus expected to be directly related to the power exponent $\eta \sim \tan\left(\frac{\pi}{2}\alpha\right)$. This means that by measuring the power-law exponent $\alpha$ independently, we should recover the value of $\eta = \frac{E''}{E'} = \frac{K''}{K'}$ that we have measured (the ratios $\frac{E''}{E'}$ and $\frac{K''}{K'}$ are equal because the proportional contribution from geometry of the cell present in both $E'$ and $E''$ cancel out)[12].

#### Supplementary material 11 | Drag coefficient

We measured the drag coefficient on the typical micropipette profile that we employed throughout this study. An oscillation of 1 µm peak-to-peak amplitude at 1Hz led to a phase lag $\varphi \sim 0.03f$ where $f$ is the frequency. During experiments the drag force is expected to be smaller, as the micropipette moves over smaller amplitudes, so this is an overestimate of the phase lag. At $f$=1Hz, $\tan\varphi \sim 0.03$, to be compared with the typical $\tan\varphi = 0.4 - 0.5$ that we measured. We subtracted a fixed estimate of this phase lag due to fluid drag, and although this could be sophisticated, we estimate an error of less that 10% in the measurement of the phase lag due to the cell only.

#### Supplementary material 12 | Effect of fast cell deformation.

We analyze raw $x_{tip}$ data by fitting a sinusoidal signal on top of a drifting accounts for the cell deformation. Intuitively if the cell deforms too fast the experimental sinusoidal signal will be drifting so fast that the fitting will become inaccurate and might introduce an artefactual elastic and viscous contribution in $x_{tip}$. We decided to test our setup and data analyzis by using a purely elastic system, where we could mimic fast cell deformation. Our strategy consisted in using a microindenter instread of a cell and to indent it with the usual setup (flexible micropipette + bead). Using a micromanipulator, we could translate this surrogate cell to mimic large cell deformation, and we check that we would measure (i) a negligible K'' and (ii) a constant K' (equal to the bending stiffness of the target microindenter) even though the target was moving. An example of rapid change in target position and resulting K' and K'' in shown in supplementary figure S7.

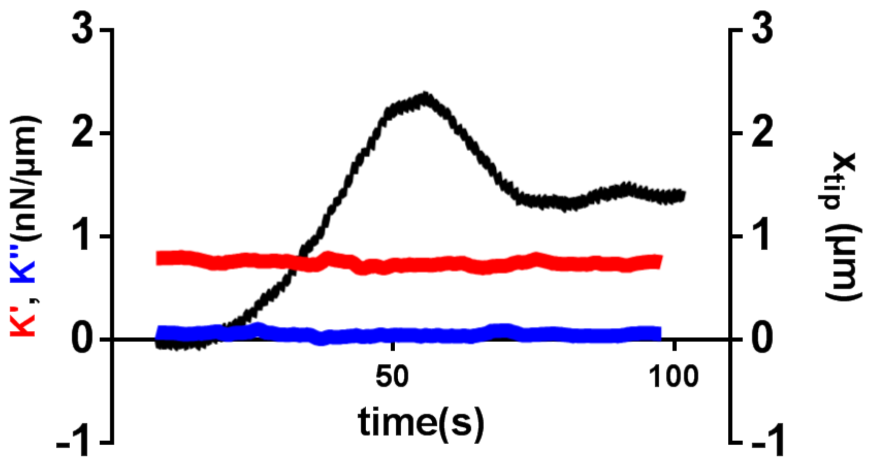


**Supplementary figure S7 |** Mimicking a fast deforming cell. The position $x_{tip}$ of the tip of a microindenter is used as a surrogate cell. The system is submitted to the protocol presented in Results section in main text. The resulting measured *K'* ansd *K''* are shown. As expected for a purely elastic object, K'' is very low. The fact that K' is constant over time shows that even with a moving target (over length and with speeds relevant to cell deformations), the data analysis leads an accurate measurement of *K'* and *K''*.
